# Refractoriness accounts for variable spike burst responses in somatosensory cortex

**DOI:** 10.1101/098210

**Authors:** Bartosz Teleńczuk, Richard Kempter, Gabriel Curio, Alain Destexhe

**Affiliations:** Unité de Neurosciences, Information & Complexité, Centre National de la Recherche Scientifique, 91198 Gif-sur-Yvette, France; Institute für Theoretische Biologie, Humboldt Universität zu Berlin, 10115 Berlin, Germany; Department of Neurology, Universitätsmedizin Charité, 12203 Berlin, Germany

## Abstract

Neurons in the primary somatosensory cortex (S1) respond to peripheral stimulation with synchronised bursts of spikes, which lock to the macroscopic 600 Hz EEG waves. The mechanism of burst generation and synchronisation in S1 is not yet understood. Using models of single-neuron responses fitted to unit recordings from macaque monkeys, we show that these synchronised bursts are the consequence of correlated synaptic inputs combined with a refractory mechanism. In the presence of noise these models reproduce also the observed trial-to-trial response variability, where individual bursts represent one of many stereotypical temporal spike patterns. When additional slower and global excitability fluctuations are introduced the single-neuron spike patterns are correlated with the population activity, as demonstrated in experimental data. The underlying biophysical mechanism of S1 responses involves thalamic inputs arriving through depressing synapses to cortical neurons in a high-conductance state. Our findings show that a simple feedforward processing of peripheral inputs could give rise to neuronal responses with non-trivial temporal and population statistics. We conclude that neural systems could use refractoriness to encode variable cortical states into stereotypical short-term spike patterns amenable to processing at neuronal time scales (tens of milliseconds).

**Significance statement:** Neurons in the hand area of the primary somatosensory cortex respond to repeated presentation of the same stimulus with variable sequences of spikes, which can be grouped into distinct temporal spike patterns. In a simplified model, we show that such spike patterns are product of synaptic inputs and intrinsic neural properties. This model can reproduce both single-neuron and population responses only when a private variability in each neuron is combined with a multiplicative gain shared over whole population, which fluctuates over trials and might represent the dynamical state of the early stages of sensory processing. This phenomenon exemplifies a general mechanism of transforming the ensemble cortical states into precise temporal spike patterns at the level of single neurons.

## 1 Introduction

Neurons usually generate highly variable responses to repeated presentations of the same stimulus. This variability might originate from thermal noise in ion channels (Chow & White, 1996; Schneidman et al., 1998), recurrent activity in the network (Destexhe et al., 2003; van Vreeswijk & Sompolinsky, 1996) or modulation of neuronal excitability (Destexhe et al., 2001; Fontanini & Katz, 2008; Faisal et al., 2008). Over recent years many results have shown that a significant fraction of this variability is shared across large populations of neurons. These shared fluctuations were attributed to the variations of incoming stimuli and modulation of excitability (Brody, 1999; Shadlen & Newsome, 1998; Goris et al., 2014; Ecker et al., 2014). However, most of these studies focused on spike-rate variations over long time scales, neglecting millisecond-range spike timing differences. Such short time scales might be especially important for neurons that fire brief bursts of spikes at a frequency reaching several hundred spikes per second separated by much longer intervals of silences (Evarts, 1964; Llinás & Jahnsen, 1982; Krahe & Gabbiani, 2004). Since the transitions between bursting and tonic firing characterised by longer interspike intervals are dynamically controlled (Swadlow & Gusev, 2001) both time scales might be relevant for neuronal processing.

Neurons in somatosensory cortices can encode their sensory inputs in the precise lengths (< 10 ms) of interspike intervals (Estebanez et al., 2012; Panzeri et al., 2001; Witham & Baker, 2015), which suggests that high firing precision is important for the reliability of stimulus encoding. In the primary somatosensory cortex (S1) of macaque, single neurons respond to peripheral stimulation with barrages of spikes elicited at sub-millisecond precision (Baker et al., 2003). However, when presented repetitively, the same stimulus produces variable responses in terms of the number of elicited spikes and the lengths of interspike intervals, which might limit the amount of information they can carry. It is, however, possible that such trial-to-trial variability represents an alternation between several classes of reliable responses, called spike patterns (Toups et al., 2012). Such spike patterns have been indeed observed in S1 (Telenczuk et al., 2011), but neither the mechanism of their generation nor their functional significance has been identified.

Here, we propose a mechanism that explains the precise patterns of single-neuron responses as an interplay between synaptic inputs and intrinsic refractory properties of the neuron (Berry & Meister, 1998; Czanner et al., 2015). To test this hypothesis, we develop simple models capturing the two processes, and we are able to fit the parameters of the models to extracellular recordings of single-unit activity in the somatosensory cortex.

## 2 Methods

### 2.1 Experimental methods

Neuronal responses were evoked in the hand representation of the primary somatosensory cortex of two awake *Maccaca mulatta* monkeys by electrical median nerve stimulation at the wrist (pulse width: 0.2 ms; repetition rate: 3 Hz; intensity: 150% motor threshold); see also Figure 1A. Single-unit activity was recorded extracellularly using a 16-channel Eckhorn drive [Thomas Recording GmbH; Giessen, Germany; Eckhorn & Thomas (1993)]. Each of the platinum/glass electrodes (electrode impedance: 1 MΩ) was advanced into cortex (area 3b) until well-isolated neurons were found with one of the electrodes. The receptive fields of the neurons were tested by means of manual tapping using a stylus.

**Figure 1:**
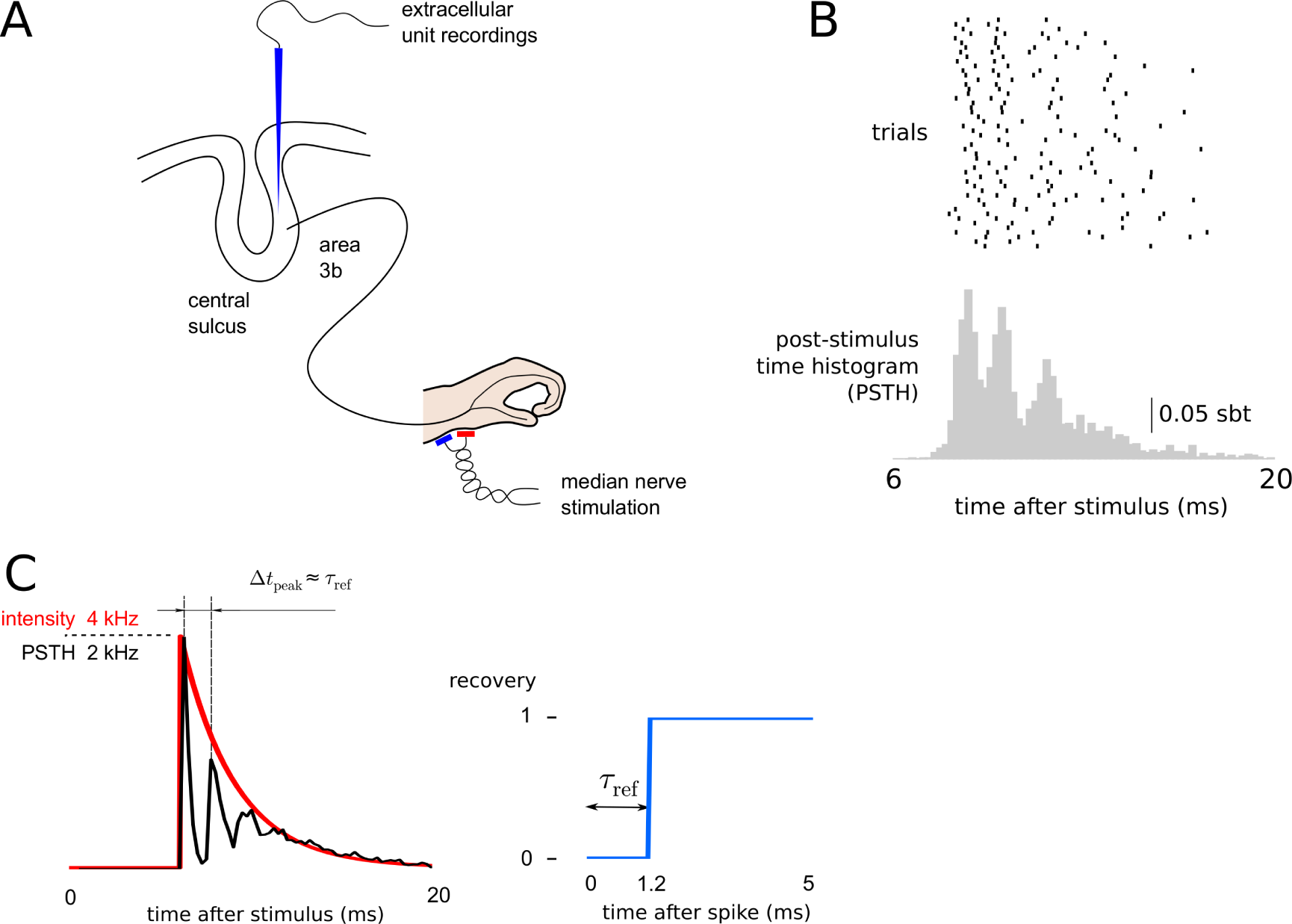
Modelling responses to median nerves stimulation of neurons recorded in primary somatosensory cortex of macaque monkeys. (**A**) Sketch of the experimental paradigm. (**B**) Raster plot of 60 sample responses of a single neuron (top) and the PSTH calculated from all 956 trials (bottom, sbt = spikes per bin per trial, bin size 0.2 ms). (**C**) Simulation of the spike-train probability model (STPM) with sample parameters: exponentially decaying intensity function (left, red line) and recovery function implementing an absolute refractory period of *τ*_ref_ = 1.2 ms (right). The simulated PSTH (left, black line) contains characteristic peaks separated by intervals approximately equal to *τ*_ref_ (left, thin vertical black lines). Note the similarity to the PSTH calculated from spikes of cortical neurons triggered by the median nerve stimulation (compare with bottom panel of B).

In addition, we recorded EEG signals from the surface of the dura (epidural EEG) with two electrodes placed in the vicinity of the micro-electrode array. The signals were then high-pass filtered (>400 Hz) to obtain the high-frequency EEG (hf-EEG).

All experimental procedures were performed according to Home Office UK (Scientific Procedures) Act 1986 regulations and institutional ethical guidelines.

### 2.2 Spike sorting

From the extracellular recording we obtained spike waveforms that were first band-pass filtered (1 kHz − 10 kHz) and then sampled with a frequency of 20 kHz. Action potentials of neurons surrounding the microelectrode were detected in the extracellular recordings by means of amplitude thresholding; the threshold was chosen manually to detect spikes whose amplitude was significantly above noise level. The wave shapes of the detected action potentials were parametrised by their amplitude, width and projection coefficients on two main principal components. The spike timings of single units were determined based on these shape features using a manual cluster cutting method that allowed for identification of clusters of arbitrary shapes (Lewicki, 1998; Hazan et al., 2006). To ensure correct clustering the procedure was performed by two operators and then checked for consistency.

In order to validate the spike discrimination, we checked the extracellular action potentials generated by a putative single neuron for the consistency of the wave shape and amplitude. Additionally, we searched for interspike intervals (ISIs) shorter than 1 ms; if such short intervals were found the clustering procedure was repeated. Spike trains with evidence of poor spike sorting (inconsistent wave shapes or ISIs < 1 ms) were excluded from subsequent analyses.

### 2.3 Spike pattern classification

From 46 neurons identified in the two monkeys we selected 17 neurons that responded with bursts of spikes. Bursting neurons were defined by responses with more than one spike for at least 4% of stimuli and a mode of the interspike interval histogram shorter than 1.8 ms (Baker et al., 2003).

Among these 17 neurons we identified neurons that also fired spontaneous bursts by counting the number of interspike intervals in the post-stimulus period (> 30 ms after the stimulus) that were shorter than 1.8 ms. In this time window the initial response dies out and baseline firing rate is re-established. Neurons that fired at least 10% of bursts in this window were labelled as spontaneous bursters.

In each neuron we summed spikes over all trials, and we identified prominent peaks in the obtained peri-stimulus time histograms (PSTH; bin width 0.2 ms, Figure 3A). As the within-burst spike composition varied from trial to trial, each trial was described with a binary string whose entries (one or zero) represented the occurrence or non-occurrence of a spike in a sequence of bins bracketing the major peaks of the overall PSTH: the borders between the bins were placed manually in the troughs of the PSTH (Figure 3A, B: vertical lines). Each binary string corresponded to one spike pattern; the length of the string equalled the total number of peaks in the PSTH.

In addition, we averaged the concomitant hf-EEG responses over trials concurring to each of the identified spike patterns of a single neuron.

### 2.4 Spike-train probability model

To reproduce the distribution of emitted spikes in a single neuron, we chose a minimal model (spike-train probability model, STPM) that could replicate the observed high variability in the cortical responses (Softky & Koch, 1993; Destexhe et al., 2001) and manifest refractoriness (decreased probability of spiking for some time after producing a spike).

We assumed that a spike emission is a random point process with the probability

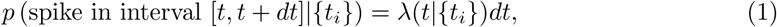

where {*t_i_*} denotes the spiking history earlier than time *t,* and *λ*(*t|*{*t_i_*}) is the conditional intensity.

The conditional intensity *λ*(*t|*{*t_i_*}) is assumed to have a Markov property, i.e., it is conditioned only on the time *t*_last_ of occurrence of the last spike at time: *λ*(*t|*{*t_i_*}) = *λ*(*t*,*t*_last_). A further assumption is that the firing-rate modulation and refractory effects are multiplicative, thus reflecting the reduction of spike probability due to, for example, inactivation of sodium channels or hyperpolarisation caused by opening of potassium channels (Berry & Meister, 1998):

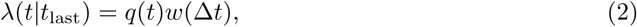

where *q*(*t*) is the intensity function, *w*(Δ*t*) is the recovery function, and Δ*t* = *t* − *t*_last_ is the time interval since the last spike.

The parameters of the model, the intensity function *q*(*t*) and the recovery function *w*(Δ*t*), are defined on a per-bin basis, and they are fitted to experimental data by means of a maximum likelihood approach. To capture fine temporal details of the neuronal responses (for example, response onset and interspike intervals) the intensity and recovery functions were defined with a short sampling interval (0.05 ms). The log-likelihood function *L*(*q*; *w|*{*t_i_*}) is obtained by log-transforming the probability function of an inhomogeneous Poisson process with the conditional intensity (2) (Dayan & Abbott, 2001; Johnson & Swami, 1983):

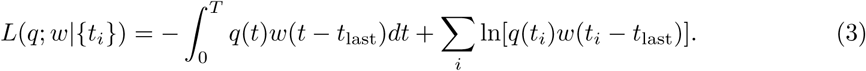

where *T* is the duration of response (*T* = 30 ms), *i* is the spike index, and *t_i_* denotes the times of occurrence of recorded spikes. The likelihood *L* of obtaining the experimental spike train *ti* is maximised with respect to the parameters *q*(*t*) and *w*(Δ*t*) by means of an iterative expectation-maximization (EM) algorithm, which guarantees that the global maximum is reached (Miller, 1985). In addition, we ensure that after 5 ms the model neuron recovers from refractoriness by setting the recovery function to unity for long intervals, i.e., we require *w*(Δ*t* > 5 ms) = 1 (for example, see Figure 2A).

**Figure 2:**
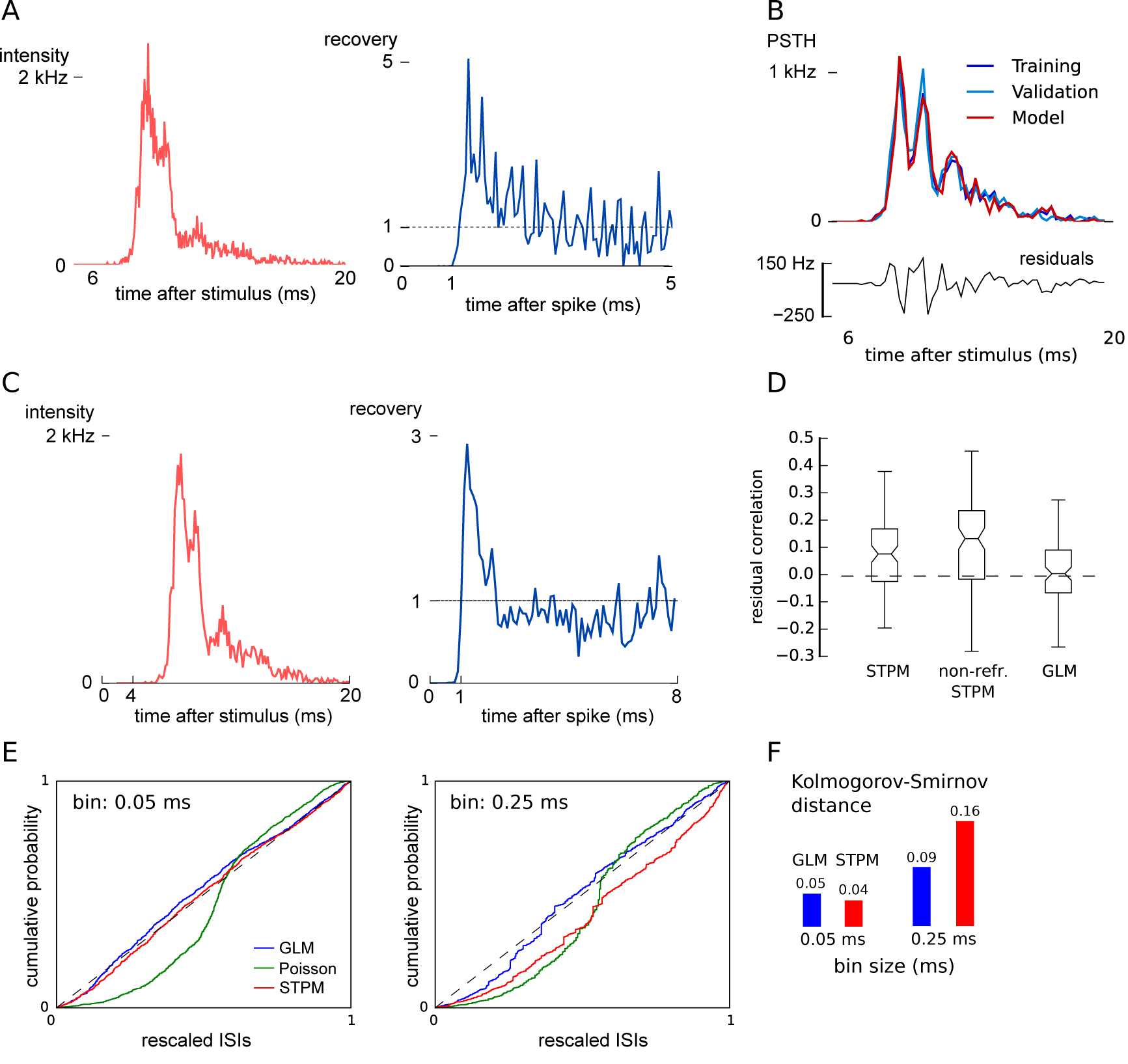
Models with refractoriness can reproduce the experimental spike trains. (**A**) Intensity function (left) and recovery (right) functions of the spike-train probability model (STPM) fitted to experimental data (an example for a single neuron). (**B**) Comparison of PSTHs of the training data (top, dark blue line), validation data (light blue line), and model data (red line). Note the overlap between the lines, which is a sign of the match between the model and both the training and validation sets. The difference between the model PSTH from the validation PSTH (model residuals, bottom) is equivalent to the intrinsic variation between the training and validation set (*F* = 1.02, *p* > 0.01, see Methods for definition). (**C**) Fitted intensity (left) and recovery functions (right) of the generalised linear model (GLM, bin size 0.25 ms). (D) Correlation coefficients between the residuals (for the STPM shown in the bottom panel in (B)) and the validation PSTH (for the STPM shown light blue in (B)) calculated for three different models: the STPM, the STPM without refractoriness (non-refr. STPM) and the GLM. The horizontal dashed line denotes the correlation coefficient between the difference of PSTH of validation and training dataset with the training dataset PSTH. (**E**) The empirical cumulative distribution of the inter-spike intervals of the experimental spike trains rescaled according to the conditional intensity function of all three fitted models (time-wrapping test). If the model perfectly reproduced the experimental inter-spike intervals the cumulative distribution should line up with the diagonal. This procedure was repeated for two different bin sizes (0.05 ms, left; and 0.25 ms, right). (**F**) The Kolmogorov-Smirnov (K-S) distance of the model (maximum divergence of the model’s cumulative distribution from diagonal in (E)) decreased with an increasing bin size in both STPM and GLM. This dependence on bin size was less pronounced for the GLM.

To study the effects of refractoriness on the modelled responses, we compared the results to the STPM without refractory period (non-refractory STPM, *w*(Δ*t*) = 1 for all Δ*t* > 0). The model is fully characterised by its intensity function *q*(*t*), which can be estimated directly from the experimental PSTH (bin width set to 0.05 ms to allow for sufficient temporal precision).

### 2.5 Generalised linear model

One limitation of the STPM is that the history effects are restricted to the last spike only. To evaluate effects evoked beyond the last spike, we considered the generalised linear model (GLM; Truccolo et al. (2005); Czanner et al. (2015)) with conditional intensity *λ*_GLM_(*t*|{*t_i_*}) of the form

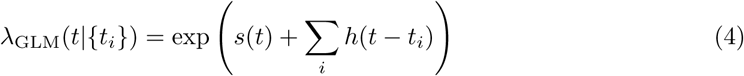

where *s*(*t*) is the driving force and *h*(*τ*) is the spike history kernel.

Note that the intensity function *q*(*t*) of the STPM can be identified with exp(*s*(*t*)), and the recovery function *w*(Δ*t*) corresponds to exp (∑_*i*_ *h*(*t* − *t_i_*)). In contrast to the STPM, in the GLM the effects of the previous spikes can extend infinitely back in time. In practice, we reduce the number of free parameters of the GLM by restricting the history horizon above which the spikes can not contribute to the responses any more; we thus set *h*(*t* > *t*_max_) = 1. The horizon *t*_max_ = 8 ms was selected to maximise the Akaike Information Criterion (AIC), which balances the goodness of fit with the number of free parameters of the model (for example, see Figure 2C).

The likelihood of the GLM is defined analogously to the spike-train probability model:

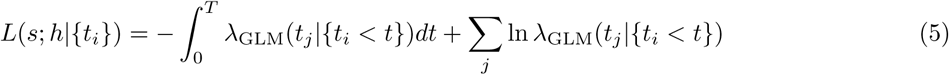

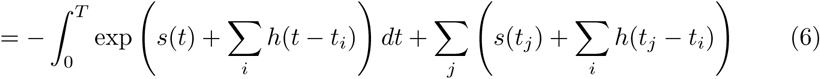

where the sums go over all spikes.

Since the log-likelihood function is a convex function of the parameters, they can be found using standard optimization techniques. In the results presented here we used the conjugate gradient optimization.

We compared the goodness-of-fit of the STPM and the GLM using the time-wrapping method (Brown et al., 2002): The inter-spike intervals in the experimental data were rescaled to account for temporal variations in firing probability. If the model perfectly reproduced the data the distribution of the rescaled inter-spike intervals would be uniform (the diagonal in Figure 2E).

### 2.6 Model validation

To validate the model, the dataset was divided into two non-overlapping subsets of equal size: a training and a validation set. The trials for each set were selected randomly from all stimulation repetitions. The parameters of the model were fitted to the training set. Based on these parameters 1000 spike trains were simulated. The goodness-of-fit was evaluated separately for two statistics *X,* that is, the PSTH (with bin size 0.2 ms) and the spike pattern distribution. For each of the two statistics, the model error was evaluated as the normalised differences between the simulated X^model^ and validations spike trains X^validate^ (cf. Rauch et al., 2003):

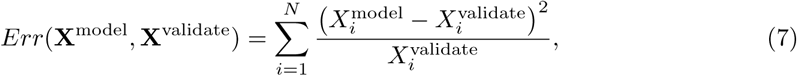

where 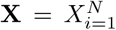 is either the PSTH or the spike pattern distribution of model (X^model^) or validation (X^validate^) set; *N* is the size of the vector and equals the number of bins (*N* = 70 for *T*=14 ms and 0.2-ms bins) or the number of identified spike patterns (*N* ≤ 16 for binary words of length less or equal to 4).

The model error *Err*(**X**^model^, **X**^validate^) was compared against the error between the training and validation sets *Err*(**X**^train^, **X**^validate^) (reference error). The significance of the difference between the model and reference errors was quantified by means of the F-test with (*N* − 1, *N* − 1) degrees of freedom (Barlow, 1989) where

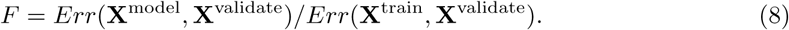

### 2.7 Serial correlations

From the responses of single neurons we identified spike triplets defined as three consecutive spikes separated by intervals shorter than 4 ms. In this analysis, to increase the number of intervals, we broadened the analysis window to 50 ms after the stimulus. Next, we calculated Pearson’s correlation between the interspike intervals (ISIs) of the first and the second spike and the second and the third spike in the triplet (*r*_data_). We compared the estimated *r*_data_ to the correlation coefficient calculated from surrogate data (*r*_model_) for the same number of trials, which were generated by the STPM model with parameters fitted to the experimental spike trains. The significance of the differences between correlation coefficients found in simulated and experimental ISIs was evaluated by means of a bootstrap test. To this end, 1000 estimates of *r*_model_ were obtained from independently simulated datasets, and the resulting coefficients were compared to *r*_data_. The *p* value was taken as the smaller of two values multiplied by 2:1) the fraction of bootstrap trials for which *r*_model_ was greater than *r*_data_ or 2) the fraction of bootstrap trails for which _*r*model_ was smaller than *r*_data_ (two-sided bootstrap test).

### 2.8 Population model

To model the relation between spike patterns of a single neuron and the response of the population, we extended the STPM to a population of uncoupled neurons receiving common inputs.

The population response was calculated from a simulated ensemble of 5000 identical neurons. The parameters of the STPM were fitted to the responses of the analyzed neuron, and these parameters were shared by all 5000 model neurons. In each trial *j* the intensity function of all neurons was modulated by a multiplicative gain factor *G_j_* that was drawn from a uniform distribution on the interval [1 − *γ*, 1 + *γ*], where 0 ≤ *γ* ≤ 1 is the strength of modulation. The intensity function in trial *j* was then *q_j_*(*t*) = *G_j_q*(*t*). From the obtained single-trial single-neuron responses the total population response was calculated by summing the binned spike responses of all neurons (bin size 0.2 ms) and subsequent band-pass filtering (400 – 1200 Hz) corresponding to the analysis of EEG data.

Next, we randomly selected a single neuron from the population and used its spikes for further analysis. We classified the spike patterns of this neuron in single trials based on the occurrence/omission of spikes in a discrete sequence of spiking “windows”. The band-pass filtered population response was then averaged over trials with respect to the type of concomitant spike pattern. This procedure, when applied to the model, reproduced the analysis that was applied to the experimental data and described above (see “Spike pattern classification”). The root mean square (RMS) amplitude of the pattern-specific average was compared with the experimentally-obtained hf-EEG related to the same spike pattern (Telenczuk et al., 2011). The similarity of the values across different spike patterns was quantified by means of Pearson’s correlation coefficient.

### 2.9 Biophysical model

In order to understand the mechanisms of burst generation, we developed a simplified single-neuron model. The model consists of a linear neuron with a spiking threshold (leaky integrateand-fire), which receives conductance-based inputs through depressing synapses (short-term synaptic depression). The membrane potential in the model follows the standard membrane equation:

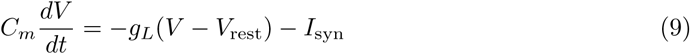

where *C_m_* is the membrane capacitance, *g_L_* is the leak conductance, *V*_rest_ is the resting potential, and *I*_syn_ are the synaptic currents. When the membrane potential reaches the threshold *V*_thr_ a spike is generated and the potential is reset to *V*_reset_ putting the cell into a hyperpolarised state. The total synaptic current in the LIF neuron is a sum of intracortical and thalamocortical currents: *I*_syn_(*t*) = *I*_Cortex_(*t*) + *I*_Th_(*t*).

The cortical synaptic currents are conductance-based inputs from *n*_inh_ inhibitory and *n*_exc_ excitatory neurons. The excitatory *g*_exc_ and inhibitory ginh synaptic conductance are a sum of contributions mediated by each spike, such that 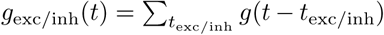. The times of the excitatory *t*_exc_ and inhibitory *t*_inh_ synaptic inputs are drawn from a homogeneous Poisson process with equal rates for excitatory and inhibitory inputs *f*_exc_ = *f*_inh_. Each spike results in a transient increase of the synaptic conductance with an exponential time course:

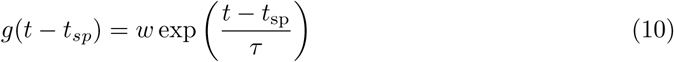

for *t* ≥ *t_sp_* and 0 otherwise. Here t_sp_ is the time of the spike, *w* is the synaptic weight and *τ* is the synaptic time constant. The reversal potentials for excitation and inhibition are *E*_exc_ and *I*_inh_, respectively. With these definitions the total current of cortical synapses is:

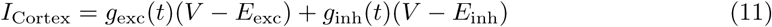

In addition to the intracortical inputs, the neuron receives excitation from nTh thalamocortical excitatory neurons. The thalamocortical neurons are silent in absence of peripheral stimulation and generate Poisson-distributed spikes 7.68 ms after the onset of the median nerve stimulus (the delay takes account of the propagation delays from periphery to the cortex). The strength of thalamocortical excitatory synapses providing the feedforward inputs to the model decays with the pre-synaptic activity following the short-term synaptic depression mechanism (Tsodyks & Markram, 1997):

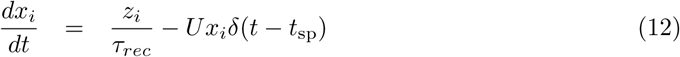

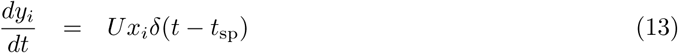

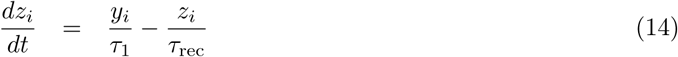

where *τ*_1_ is the decay constant of synaptic conductance, *τ*_rec_ is the recovery time from synaptic depression and *U* describes the fraction of available resources used by each presynpatic spike.

The postsynaptic current due to the thalamocortical inputs is then:

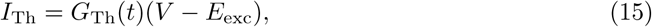

where the total conductance due to thalamic inputs is given by:

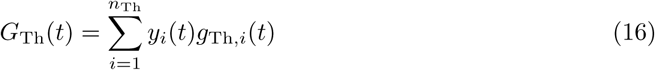

*g*_Th,*i*_ stands for conductance of a single synapse and *y_i_* its efficiency. The contribution of a single presynaptic spike to postsynaptic conductance is assumed to be the same as for an excitatory cortical synapse (i.e. *g*_exc_(*t*)).

Eight parameters of the model were adjusted to reproduce the experimental PSTH: weights of excitatory (*w*_exc_), inhibitory (*w*_inh_) and thalamocortical (*w*_Th_) synaptic inputs, excitatory synaptic time constant (*τ*_exc_), firing rates of cortical (*f*_exc_) and thalamocortical (*f*_Th_) pre-synaptic neurons, number of thalamocortical synapses (*n*_Th_), and use of synaptic resources by thalamocortical synapse release (*U*). Other parameters were fixed to values found in the literature. The values of other parameters are given in Table 1.

**Table 1:**
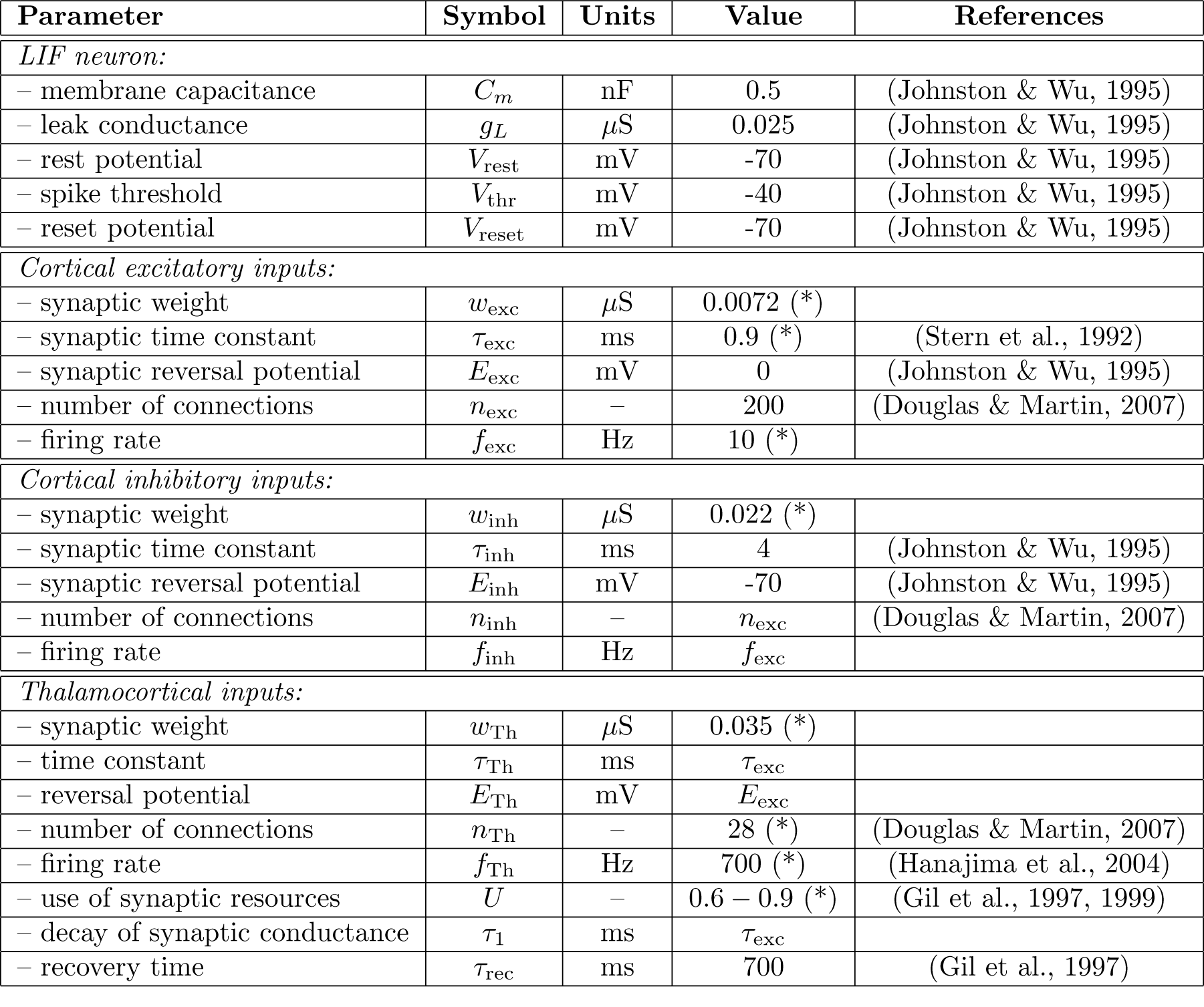
List of parameters used in the leaky integrate-and-fire model. “Value” column indicates typical parameter values or ranges found in the literature (where available). (*) denotes the parameters which were adjusted to fit the experimental data.

| Parameter | Symbol | Units | Value | References |
| --- | --- | --- | --- | --- |
| *LIF neuron:* |  |  |  |  |
| – membrane capacitance | $C_m$ | nF | 0.5 | (Johnston & Wu, 1995) |
| – leak conductance | $g_L$ | $\mu S$ | 0.025 | (Johnston & Wu, 1995) |
| – rest potential | $V_{\text{rest}}$ | mV | -70 | (Johnston & Wu, 1995) |
| – spike threshold | $V_{\text{thr}}$ | mV | -40 | (Johnston & Wu, 1995) |
| – reset potential | $V_{\text{reset}}$ | mV | -70 | (Johnston & Wu, 1995) |
| *Cortical excitatory inputs:* |  |  |  |  |
| – synaptic weight | $w_{\text{exc}}$ | $\mu S$ | 0.0072 (*) | |
| – synaptic time constant | $\tau_{\text{exc}}$ | ms | 0.9 (*) | (Stern et al., 1992) |
| – synaptic reversal potential | $E_{\text{exc}}$ | mV | 0 | (Johnston & Wu, 1995) |
| – number of connections | $n_{\text{exc}}$ | – | 200 | (Douglas & Martin, 2007) |
| – firing rate | $f_{\text{exc}}$ | Hz | 10 (*) | |
| *Cortical inhibitory inputs:* |  |  |  |  |
| – synaptic weight | $w_{\text{inh}}$ | $\mu S$ | 0.022 (*) | |
| – synaptic time constant | $\tau_{\text{inh}}$ | ms | 4 | (Johnston & Wu, 1995) |
| – synaptic reversal potential | $E_{\text{inh}}$ | mV | -70 | (Johnston & Wu, 1995) |
| – number of connections | $n_{\text{inh}}$ | – | $n_{\text{exc}}$ | (Douglas & Martin, 2007) |
| – firing rate | $f_{\text{inh}}$ | Hz | $f_{\text{exc}}$ | |
| *Thalamocortical inputs:* |  |  |  |  |
| – synaptic weight | $w_{\text{Th}}$ | $\mu S$ | 0.035 (*) | |
| – time constant | $\tau_{\text{Th}}$ | ms | $\tau_{\text{exc}}$ | |
| – reversal potential | $E_{\text{Th}}$ | mV | $E_{\text{exc}}$ | |
| – number of connections | $n_{\text{Th}}$ | – | 28 (*) | (Douglas & Martin, 2007) |
| – firing rate | $f_{\text{Th}}$ | Hz | 700 (*) | (Hanajima et al., 2004) |
| – use of synaptic resources | $U$ | – | 0.6 – 0.9 (*) | (Gil et al., 1997, 1999) |
| – decay of synaptic conductance | $\tau_1$ | ms | $\tau_{\text{exc}}$ | |
| – recovery time | $\tau_{\text{rec}}$ | ms | 700 | (Gil et al., 1997) |

## 3 Results

Neurons in area 3b of macaque monkeys show brief (< 10 ms) bursts of activity in response to stereotypical electrical stimulation of the median nerve (0.2 ms pulse, 1.5 time motor threshold applied transcutaneously to the median nerve); see also Figure 1A. In a dataset of 46 neurons recorded extracellularly using movable platinum-glass electrodes (Eckhorn drive, Thomas Recordings) we found 17 neurons that responded with burst of spikes (defined as trains of two or more spikes with interspike intervals shorter than 1.8 ms). When averaged over several repetitions of the stimulation the responses gave rise to a post-stimulus time histogram (PSTH) with prominent peaks coincident with the within-burst spikes (Figure 1B). The appearance of such PSTH peaks points to the precision of the burst timing with respect to the onset of the stimulus.

Some of these bursting neurons also elicited spikes in absence of median nerve stimulations (5 neurons fired at least 10% of bursts in the window [30, 300] ms after the stimulus). The evoked and spontaneous bursts differed slightly with respect to mode of the within-burst interval distribution [evoked: 1.82 (1.71) ms; spontaneous: 1.32 (0.41) ms, mean (standard deviation) across neurons] and burst length [evoked: 2.76 (1.26) spikes per burst; spontaneous: 2.18 (0.39) spikes per burst], but these differences were not statistically significant (t-test, *p* > 0.01).

To understand the mechanisms underlying bursting of neurons in the primary somatosensory cortex, we propose a phenomenological model of the single-neuron response to the median nerve stimulation. The model is based on two experimental observations: 1) Upon presentation of strong sensory stimuli, layer IV cortical neurons are bombarded with intense and coincident synaptic inputs from thalamocortical neurons (Bruno & Sakmann, 2006; Hanajima et al., 2004; Swadlow & Gusev, 2001; Gil et al., 1999; Cruikshank et al., 2007). 2) After emitting a spike, neurons are refractory, which limits their maximum firing rate (Berry & Meister, 1998; Gray, 1967; Kara et al., 2000). In order to illustrate the effects of these two phenomena on neuronal responses, we simulated a probabilistic model (the spike-train probability model, STPM, see Methods) with an exponentially decaying intensity function and an absolute refractory period *τ*_ref_ = 1.2 ms (Figure 1C). The PSTH of the simulated spike responses qualitatively reproduces main features of the PSTH obtained from experimental data. Specifically, the absolute refractory period leads to an appearance of multiple peaks in the PSTH (three peaks visible in Figure 1C: at 6 ms, 7.8 ms, and 9.5 ms after the stimulus) separated by deep troughs corresponding to periods of quiescence during which the neuron is refractory. The presence of such peaks and troughs in the trial-averaged PSTH is possible because the responses of the neuron are reliable across trials. The first peak of the PSTH reflects the initial spike triggered by the sharp transient of the intensity function (Figure 1C, red line). This initial response is highly reliable, giving rise to the narrowest and tallest PSTH peak (half-amplitude width: 0.75 ms; peak-to-trough amplitude: 1773 spikes/s in Figure 1C, black line). The refractory state following the first spike leads to a pronounced decrease of firing probability and gives rise to the deep PSTH trough following the initial PSTH peak. The subsequent PSTH peaks become wider and are of smaller amplitude due to the gradual decay of the intensity function (second peak: 1.25 ms, 1002 spikes/s; third peak: 1.25 ms, 290 spikes/s). The PSTH obtained from this simulation is qualitatively similar to cortical burst responses triggered by peripheral nerve stimulation (compare Figure 1C, left with Figure 1B, bottom).

### Refractoriness explains the intra-burst intervals

We demonstrated that the STPM with a decaying intensity function and an absolute refractory period can produce a PSTH that agrees qualitatively with the responses of neurons in primary somatosensory cortex of macaques. In order to test whether the STPM can also quantitatively reproduce the fine details of neuronal responses recorded in vivo, we inferred the intensity and recovery functions directly from the data. The two functions were defined on per-bin basis and were treated as the free parameters of the model. These parameters were then fitted to the experimental spike trains using a convex optimisation technique guaranteeing the identification of the most optimal model (see Methods, Figure 2A).

The fitted intensity function peaks shortly after the stimulus onset (< 10 ms) and decays back to baseline when the burst is terminated (Figure 2A, left). The intensity function still contains three distinct peaks, but they are less prominent compared to the peaks in the PSTH (Figure 1B). This smoothing can be attributed to the decoupling of synaptic inputs, which are represented by the intensity function, from the refractoriness, which is represented by the recovery function (Figure 2A, right). Although the maximum of the intensity function is much above the rate at which individual neurons can fire spikes, the refractoriness limits the firing rate of the model neuron. In agreement with the properties of biological neurons, the fitted recovery function is equal to 0 for the first 1 ms after emitting a spike (absolute refractory period), but after a few milliseconds fully recovers from the refractoriness returning to the rest state (*w*(*t*) ≈ 1). Interestingly, immediately after the absolute refractory period the recovery function over-shoots for about 1 ms, largely exceeding the rest value. The fast (≪ 1 ms) fluctuations following this over-shoot represent statistical noise due to the finite size of the data set. Altogether, the parameters of the STPM disentangle the synaptic inputs from the refractory effects.

### The STPM provides a parsimonious description of bursting in S1 cortex

The simulated peri-stimulus time histogram (Figure 2B, red line) matches closely the one obtained from the experimental data (Figure 2B, dark blue line). In order to demonstrate that this good match is not a result of an over-fitting, we performed cross-validation. First, the data set was divided randomly into two subsets: training data and validation data. The model was fitted only to the first subset, and then the results of the simulation were validated on the second (Figure 2B, light blue line). We found that the difference of the fitted PSTH from the validation set was of the same magnitude as the variation within the dataset (see Methods, F-test, *p* > 0.01). This test indicated that the model optimally captured the features of both training and validation set without considerable over-fitting.

The parameters of the model were fitted to each of the 17 neurons yielding similar results. Importantly, an application of the cross-validation procedure revealed that in 12 out of the 17 neurons the PSTH simulated with the model was not significantly different from the PSTH calculated from the recorded spike trains (*F* = 0.65 −1.59, *p* > 0.01). In the remaining 5 neurons the modelled PSTH deviated significantly from the validation PSTH (*F* = 2.33 − 4.88, *p* < 0.01, F-test). This sub-population of neurons may have differing firing properties that would need more sophisticated models (implementing, for example, bursting mechanisms; see Discussion).

To further analyse the cases in which the simulated spike trains differed from the data, we subtracted the model PSTH from the validation data PSTH (Figure 2B, bottom). The resulting residuals still contained fluctuations aligned to the peaks of the PSTH. This indicated that the model does not fully capture the shape of the PSTH. Indeed, the correlation coefficient between the residuals and the PSTH of the validation set was significantly positive (bootstrapped 95% confidence intervals, Figure 2D) meaning that the residuals contain some remnants of the averaged neuron response. Altogether, these results show that the STPM model is sufficient for describing trial-averaged responses in the majority of recorded neurons.

### Poisson-like variability explains the occurrence of temporal spike patterns in repeated trials

Having shown that the interplay between the intensity and recovery functions of the STPM can account for a large part of the trial-averaged response of a single neuron, we tested whether the model can also explain the trial-to-trial variability of the spiking of cortical neurons.

In order to quantify the trial-to-trial variability of neuronal responses, we sorted single-trial spike trains according to the occurrence of spikes in pre-defined temporal windows (Figure 3A-C). Each spike train was assigned a binary word based on occupancy of preferred firing windows the borders of which were aligned to the troughs of the PSTH (windows labelled ‘x’, ‘y’ and ‘z’ in Figure 3A). As explained above, these troughs reflect the periods of quiescence due to the refractoriness of the neuron. When the single-trial spike trains were re-ordered according to the associated binary word, we could distinguish between several patterns of activity (spike patterns). In most trials the neuron fired in all three windows (triplet, 111) or only the first two (doublet, 110), where the input was the strongest (compare with Figure 2A), but also doublets with other combinations of spikes and silences were common (the spike pattern frequency distribution is shown in Figure 3C). For example, the doublet 101 corresponds to trials in which the neuron fired at the onset of the stimulation (window ‘x’, Figure 3A), then remained silent during the second window (‘y’) and fired again in the third window (‘z’); the omission of the spike in the window ‘y’ is the consequence firing late in the window ‘x’ (see the corresponding line in the raster plot of Figure 3B), so that the neuron is refractory during the window ‘y’. In other neurons the number of discrete firing windows (determined by the number of PSTH peaks) ranged from two to four, and a similar distribution of spike patterns was obtained. The appearance of such spike patterns can be attributed to the chance phenomena (Poisson firing) and their interplay with the structured input and refractoriness.

**Figure 3:**
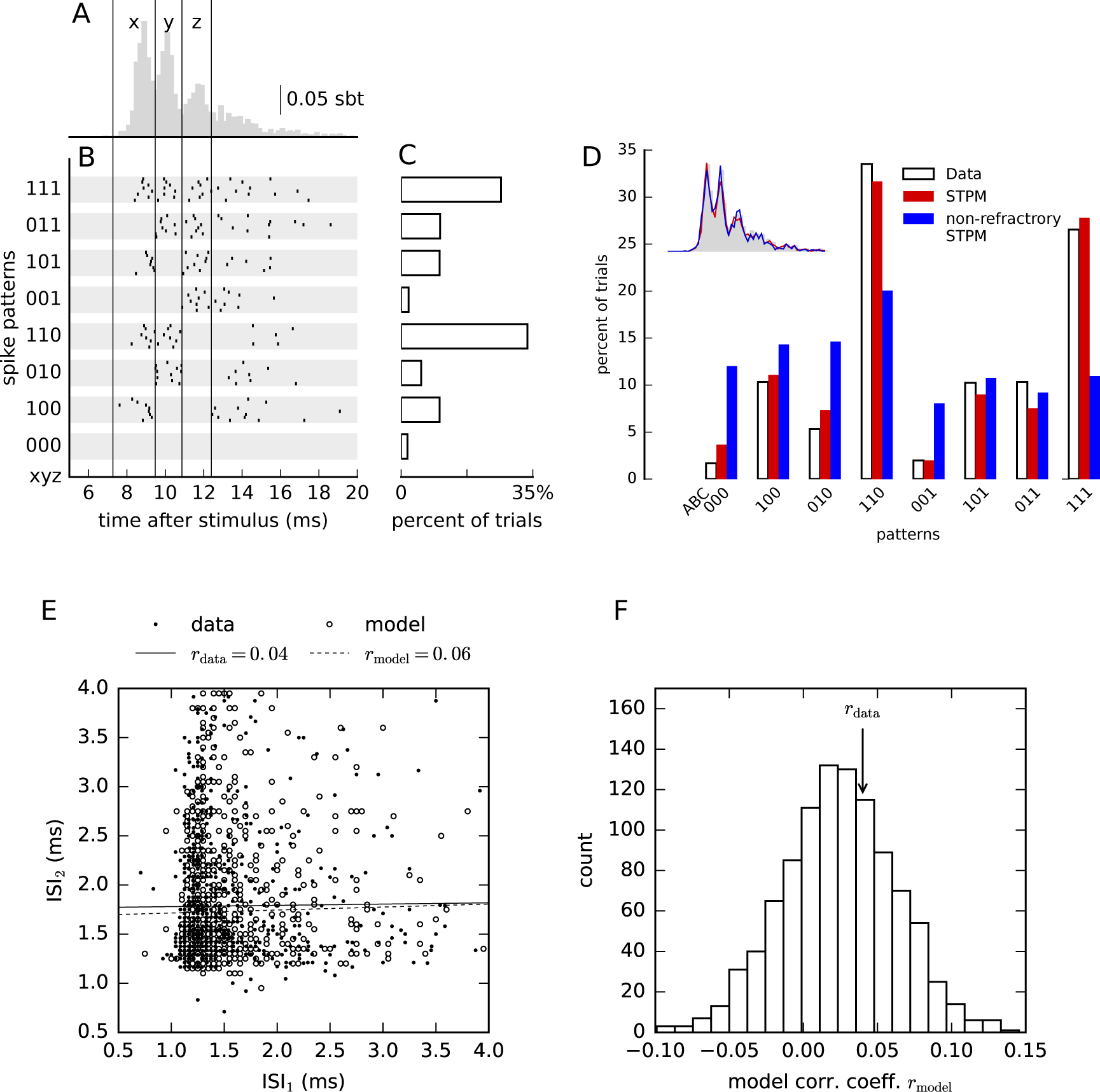
The STPM explains trial-to-trial variability of the data. (**A**) Single-neuron responses averaged over all trials (PSTH, same as in Figure 1B) reveal that spikes occur preferentially at discrete latencies (delimited by vertical lines and indexed by ‘x’ for the first peak, ‘y’ for the second peak, and ‘z’ for the third peak). (**B**) In single trials multiple spikes are elicited in diverse combinations of preferred latencies resulting in significant trial-to-trial response variability. Spike combinations are classified into spike patterns: The time axis was first divided into three windows aligned to the peaks of the PSTH. Each trial was then assigned a binary string (spike pattern ‘xyz’, from ‘000’ to ‘111’), where 1 represents the occurrence and 0 the absence of a spike in a window. Spike timings of eight representative sample responses assigned to each pattern are shown as raster plots. (**C**) Frequency at which the spike patterns occurred over repeated trials for the neuron in (A). (**D**) Firing pattern distribution obtained from the data (white bars, same as C), the STPM (red bars) and the non-refractory STPM (blue bars). The firing rate of the Poisson model was estimated by a PSTH with bin size 0.05 ms. Inset compares the PSTHs obtained from each model (color-coded like the bars in main panel). (**E**) Scatter plot of two consecutive interspike intervals (ISIs) within spike triplets calculated from the experimental data (filled circles) and responses simulated with STPM (empty circles). Serial correlations (Pearson’s correlation coefficient) found in the experimental intervals (r_data_) differ only slightly from the respective correlations predicted by the STPM (r_model_, see values in the legend, solid and dashed lines represent the best linear fit to the experimental and model data, respectively). (**F**) Repeated Monte-Carlo simulations (*n* = 1000) of the STPM fitted to experimental data provide the distribution of serial correlations consistent with the STPM (empty bars); the serial correlations estimated directly from experimental data (vertical arrow, r_data_) are likely to be drawn from the same distribution (two-sided bootstrap test, *p* = 0.81).

We found that the distribution of spike patterns in the experimental data was similar to the distribution obtained from the STPM (Figure 3D). In contrast, when the recovery function was constrained to 1 for all bins and the intensity function estimated from the data (non-refractory STPM) some spike patterns appeared at frequencies much different from the experimental data (for example, spike patterns 010, 110, 001 and 111 in Figure 3D), despite the fact that the overall PSTHs were almost identical (Figure 3D, inset). The differences of spike pattern frequencies can thus be understood as the effect of refractoriness; without it the probabilities of firing in each window are independent of the occurrence of spikes in the previous windows, in which case the frequency of a spike pattern can be directly predicted from the trial-averaged response (PSTH).

To quantify the similarity between the experimental and modelled spike patterns, we used a cross-validated chi-square statistics (see equation (7) in the Methods). In 12 of 17 examined neurons the spike-pattern distribution of the STPM was similar to the experimental distribution, and for 5 cells they were significantly different (F-test, *p* < 0.01); in 2 of these 5 neurons the PSTH was not accurately predicted by the STPM precluding the possibility of predicting the trial-to-trial variations. In the remaining 3 neurons there were substantial differences in the frequency of selected spike patterns, which might reflect the mis-estimation of the recovery function. Overall, these results show that in most neurons the STPM with time-dependent inputs and refractoriness can account not only for the trial-averaged but also the trial-to-trial variability of responses to somatosensory stimulation.

### Within-burst intervals manifest significant correlations

Next, we investigated whether the correlations between consecutive interspike intervals (serial correlations) may play a role in the generation of spike patterns. The STPM predicts that the response should be fully determined by the current input and the interval since the last spike. However, the calculation of the serial correlations in the experimental data obtained from S1 showed that two consecutive interspike intervals are not independent (Figure 3E). Since significant serial correlations might be induced by the firing-rate variations alone, we compared the experimental serial correlations with the ones obtained with the STPM, which does not assume any correlations between interspike intervals. In the example shown in Figure 3E the serial correlations are indeed accounted for by the STPM model meaning that the spiking history prior to the last spike does not affect the response.

In 12 out of 17 neurons the experimental and model serial correlations were not significantly different (two-sided bootstrap test, *p* > 0.01, Figure 3F) confirming that for most neurons the spiking memory did not extend over the last spike. In 3 neurons the coefficient could not be determined because of a low number of triplets identified in responses. In 2 neurons the correlation coefficients were larger in the data than in the fitted STPM model (bootstrap test, *p* < 0.01).

We also compared the STPM with a generalised linear model (GLM, Figure 2C), which can account for spiking history extending over the last spike. The GLM showed a similar power in explaining both the average PSTH compared to the STPM (t-test, *p* < 0.01, Figure 2D, right box). However, it allowed for using larger bins without significant loss of goodness-of-fit (Figure 2E, F). Finally, the introduction of spike history effects extending over multiple preceding spikes did not explain the significant serial correlations in every neuron. The GLM could account for the measured serial correlations in 13 out of 17 neurons. Overall, these results show that refractoriness is sufficient to explain the statistics in the within burst intervals obtained in most recorded neurons.

### Trial-to-trial input variations induce significant serial correlations

The significant serial correlations found in two neurons could result from variability of the inputs that they receive. Although the peripheral stimulation of the median nerve used to evoke the somatosensory responses was well controlled over the duration of the recording, it is possible that the effective input to the cortex was modulated at the early stages of somatosensory pathway (cuneate nucleus, thalamus) and by on-going activity in the cortex. On the other hand, the STPM was fitted under the assumption that the inputs and model parameters do no change in time, i.e., that they are stationary.

To test the effects of trial-to-trial variability on the estimated STPM parameters and the serial correlations, we simulated spike trains from the STPM with a step-like recovery function and an exponentially decaying intensity function (Figure 4B and C, dashed lines). In addition, in each trial we modulated the amplitude of the intensity function by a multiplicative gain, G, which was randomly drawn from uniform distribution on the interval [0.2, 1.8] (Figure 4A). Next, we fitted the simulated surrogate data with an STPM assuming that the intensity function was fixed and that the trial-to-trial variability resulted solely from the probabilistic nature of the model. The fitted intensity function (Figure 4B, red line) reflected the rapid onset and slower decay of the input after the stimulus, but its trace deviated from the “ground-truth” intensity function used in the simulation (compare the solid red and dashed gray curves in Figure 4B). Importantly, the intensity function contained small ripples akin to the ones visible in the intensity function fitted to experimental data (Figure 2A, left). Similarly, the fitted recovery function did not capture the step-like transition from refractoriness to baseline, but it manifested a prominent overshoot following the absolute refractory period and slower decay to baseline (Figure 4C); such time-dependence was reminiscent of the shape of recovery function estimated from the data (Figure 2A, right).

**Figure 4:**
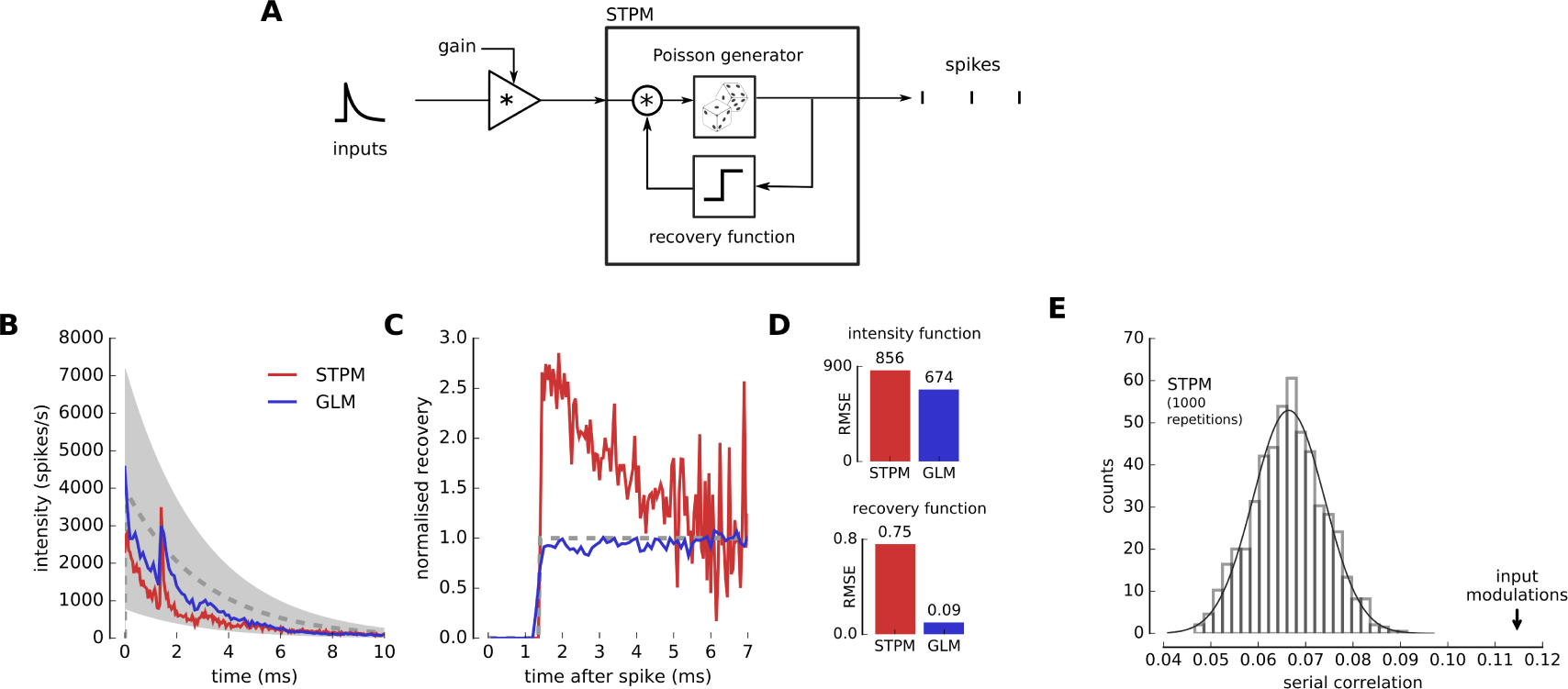
Input modulation may explain deviations of spike train statistics from the STPM and GLM. (**A**) The STPM was extended by including a multiplicative gain factor, which acts on the input function. The gain factor was randomly selected from a uniform distribution [0.2, 1.8] in each trial. The model was simulated with an exponentially decaying intensity function (dashed line in (B); maximum amplitude, 4000 spikes/s, time constant, 3 ms) and a step recovery function (dashed line in (C); refractory period, 1.4 ms). (**B**) Intensity function of the STPM (red) and GLM (blue) fitted to the simulated spike trains. The intensity function manifests deviations from the real intensity function used in the simulation (dashed line; gray-shaded area corresponds to the amplitude range of intensity function taking into account the gain factor). (**C**) Recovery function of the STPM (red) and GLM (blue) fitted to the simulated spike trains. The STPM-estimated recovery function displays a characteristic overshoot soon after end of the absolute refractory period (1.4 ms; dashed line – real recovery function underlying the spike trains). (**D**) Root-mean-square error of the intensity and recovery functions estimated with the STPM and GLM. (E) The serial correlation of the model with gain modulation (arrow) is significantly larger (p *<* 0.01) than predicted in absence of modulation (bar plot – histogram of 1000 serial correlation coefficient obtained from Monte Carlo simulations of the STPM with the intensity and recovery functions shown in (B) and (C, red line)).

We also studied the effects of the gain modulation on the GLM. The intensity function estimated with this model still contained fluctuations absent in the function used for simulation, but their amplitude was reduced compared to the STPM. A greater improvement was observed in the GLM estimate of the recovery function, which approximated well the real function without a visible overshoot. Overall, both STPM and GLM mis-estimated some model parameters in presence of trial-to-trial variation, but we found that the GLM was more robust (Figure 4D).

Finally, we estimated the serial correlation between the interspike intervals in presence of the input modulation. We found that the serial correlation was significantly larger compared to the spike trains simulated with the STPM with no trial-to-trial variations (Figure 4E). This result shows that positive serial correlations can be obtained when neuronal responses vary from trial to trial reflecting changing inputs or excitability of the neuron. Since in our analysis in Figures 3E and F we compared experimental serial correlations to the ones obtained from the STPM, which does not account for the input variability, our estimate of serial correlations could reflect input modulation.

In summary, we show that the trial-to-trial variations of the input can explain several aspects of the STPM fitted to experimental data, in particular the ripples in the fitted intensity functions, overshoot following the refractoriness in the recovery function, and significant correlations between consecutive interspike intervals.

### Trial-to-trial input variations induce correlations between single-neuron and population responses

Simultaneous recordings of single-neuron spike patterns and macroscopic EEG signals recorded from the surface of dura (high-frequency, > 400 Hz, epidural EEG) have shown that the spike patterns are not private to each neuron but that they are coordinated across a population of neurons responding to peripheral stimulation (Telenczuk et al., 2011). Such a coordination could possibly by achieved with a millisecond range-synchronisation of the neurons, but the mechanisms of such a synchronisation are not clear. Alternatively, it could be produced by the shared modulation of inputs or excitability. In order to test the latter hypothesis, we applied our probabilistic single-neuron model, the STPM, to a population of neurons receiving common gain modulation (Figure 5A).

**Figure 5:**
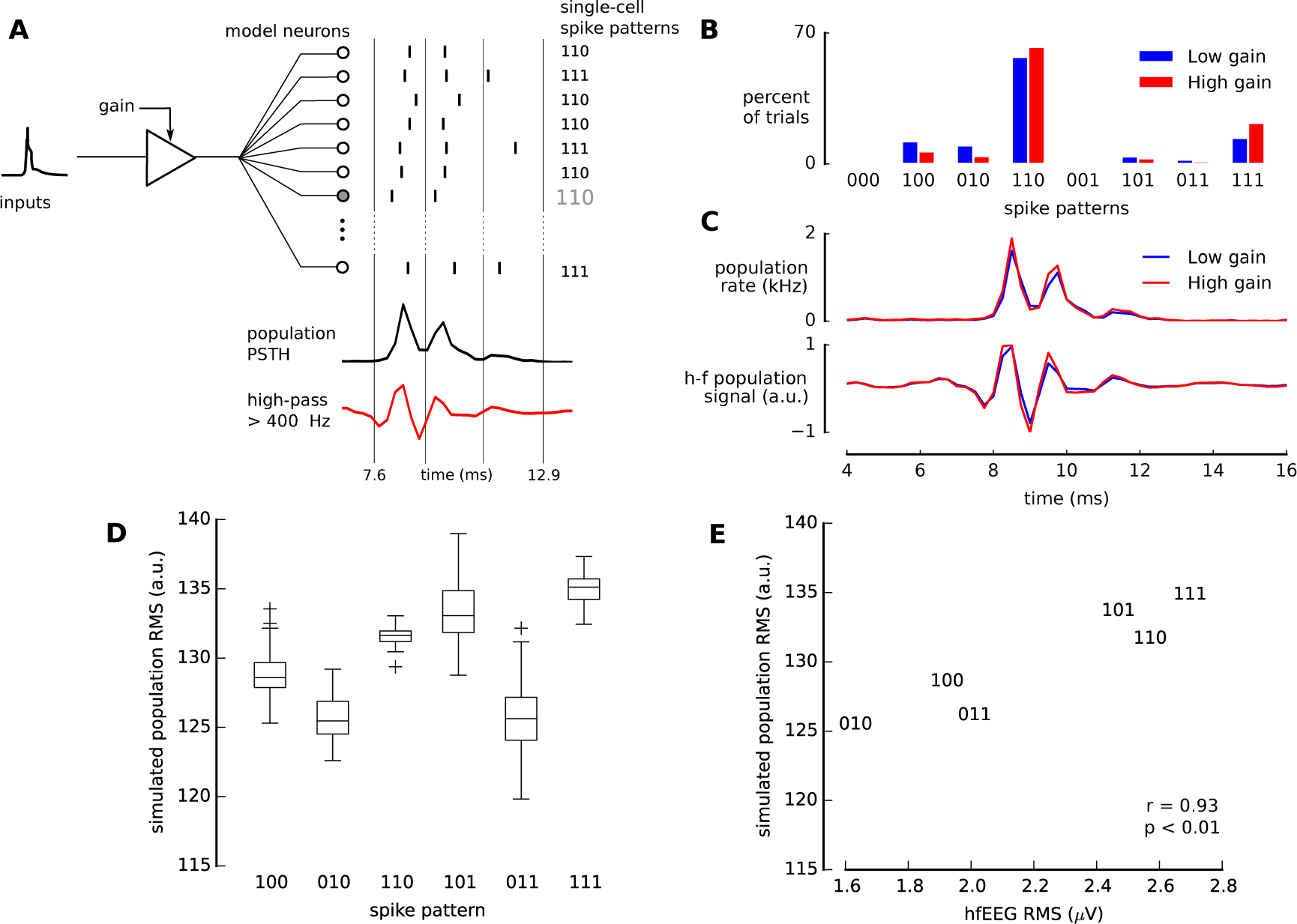
Coordination of spike patterns in the population. (**A**) Simulation of 5000 identical units described by the STPM (Figure 2C) with gain modulation of the strength *γ* = 0.2. From the simulated spike trains of all neurons the population PSTH was calculated and then high-pass filtered to obtain an estimate of the high-frequency EEG population response. (**B**) Distributions of spike patterns of a single neuron in 1000 repetitions of the simulation with a low (0.8, blue) and a high (1.2, red) gain. (**C**) The population PSTH before (top panel) and after high-pass filtering (bottom panel) varies with the gain (blue: 0.8, red: 1.2). (**D**) Single-neuron spike pattern and root-mean-square (RMS) amplitude of the high-pass filtered population PSTH are correlated because both the spike pattern and the PSTH depend on the gain. (**E**) The simulated population RMS amplitudes correlate with experimental hf-EEG RMS related to the same pattern (hf-EEG RMS) (Telenczuk et al., 2011).

As before, we assumed that the gain varies from trial to trial due to fluctuations in excitability, synaptic strength, or background activity. In order to investigate the effect of the gain factor on the population response, we simulated 5000 identical, statistically independent model neurons with the parameters estimated from the experimental data. We found that the frequencies of individual spike patterns depended on the value of the gain factor: Some patterns (for example “100”) occur more frequently at low gain (G=0.8), while others (for example “110”) tend to occur more often at high gain (G=1.2, Figure 5B). Concurrently, the amplitude of the binned spike trains averaged across neurons (population PSTH) increased with the gain (Figure 5C).

The concurrent dependence of population PSTH and single-neuron spike pattern distribution on the gain factor may also explain the correlation between single-neuron responses and macroscopic population activity found in experimental data. Spike patterns that are more frequent at low gain coincide predominately with a low-amplitude population PSTH whereas spike patterns elicited more frequently at high input gain, on average, coincide more often with a high-amplitude population PSTH. Consequently, the amplitude of the population PSTH might co-vary with single-neuron spike patterns. In particular, we found that the root-mean-square amplitude of the high-pass filtered PSTH (> 400 Hz) depends on the spike pattern used for grouping the trials (Figure 5D).

To test whether gain modulation could explain the experimental results, we simulated the STPM model with trial-varying gain factor (see Methods) and compared the spike-pattern-specific high-frequency EEG (hf-EEG) amplitude calculated from experimental data with the simulated population response. We found that already for a modest level of the gain modulation (modulation strength *γ*=0.2) the root-mean-square amplitudes of the experimental hf-EEG and high-pass-filtered population PSTH of the model were strongly correlated (Figure 5E, an example for one neuron, Pearson’s *r* = 0.93).

We found a positive correlation coefficient in 12 of 16 neurons that produced at least 3 different patterns. This fraction is significantly above the chance level expected from uncorrelated quantities (two-sided binomial test, p<0.05). Thus, we conclude that gain modulation can introduce correlations between the single-neuron spike patterns and macroscopic population responses.

### Spike patterns emerge as input-driven phenomena in a simplified biophysical model of a cortical neuron

The probabilistic models presented so far are abstract, and their parameters (intensity and recovery functions) can not be linked directly to biophysical properties of a neuron. To interpret the generation of spike patterns mechanistically, we developed a simplified biophysical model of a cortical neuron based on the leaky integrate-and-fire (LIF) model. Although this model does not reproduce faithfully all biological properties of realistic neurons, it captures their integration and spike generation properties, which are essential to the responses analysed here. We simulated the neuron with two types of synaptic inputs – tonic excitatory and inhibitory inputs, and phasic thalamic excitatory inputs representing the barrage of action potentials triggered by peripheral stimulation.

In absence of thalamic inputs the model neuron elicits only few spikes due to spontaneous threshold crossings. However, in the model the median nerve stimulation is assumed to activate the thalamocortical fibers (28 synapses per cortical neuron), which then fire randomly according to a Poisson distribution with the rate of 700 spikes per second. These massive inputs trigger excitatory post-synaptic currents bringing the membrane quickly to the threshold. This results in a series of spike emissions accompanied by rapid successions of membrane de- and re-polarisations (Figure 6A).

**Figure 6:**
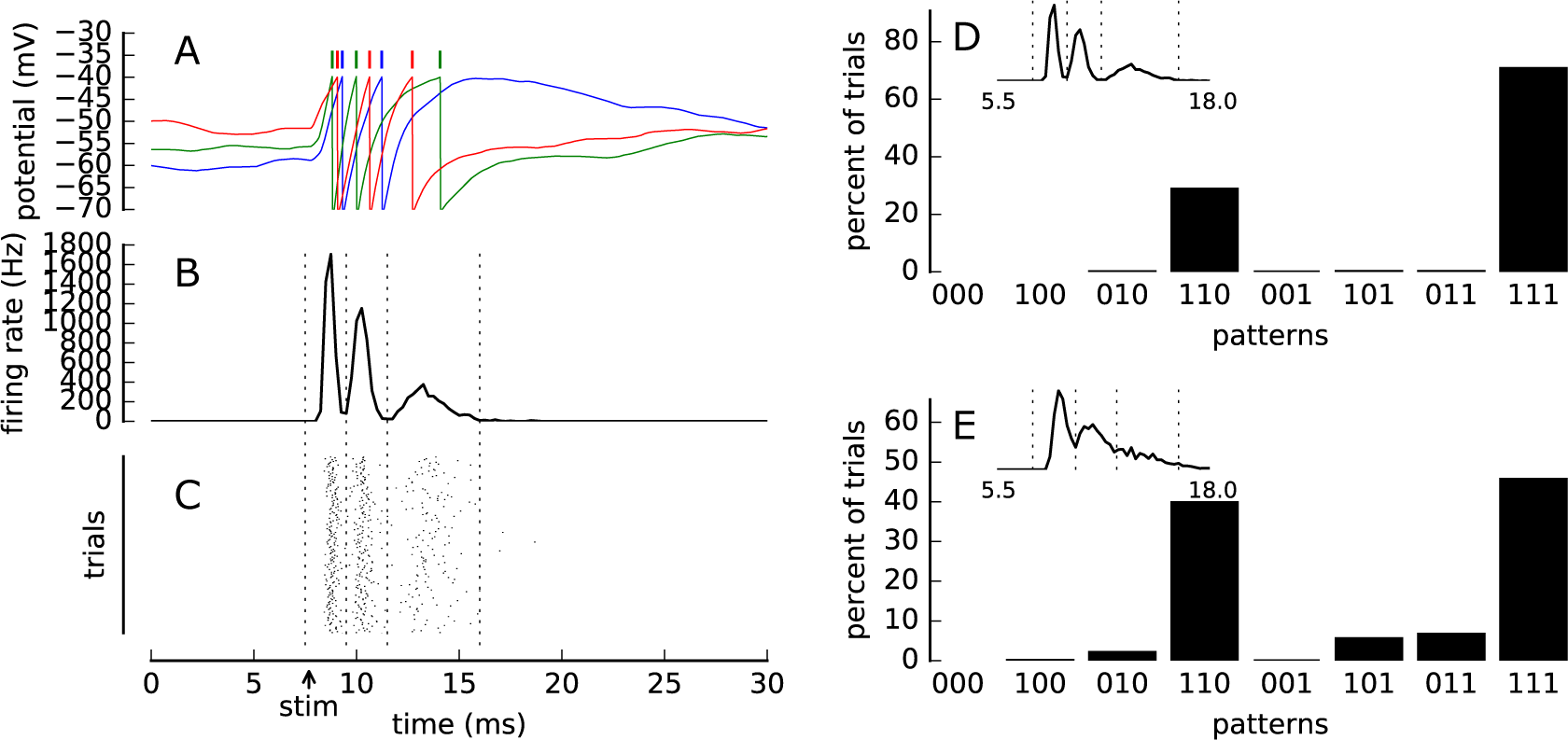
A leaky integrate-and-fire (LIF) model produces variable spike patterns. (**A**) Sample traces of the membrane potential *V_m_*(*t*) of a leaky integrate-and-fire model (see Methods for details) for three repetitions of the simulation. The ticks mark the threshold crossings, which lead to spike emission (color matched to the color of *V_m_* trace). (**B**) Post-stimulus time histogram (firing rate) of spike trains obtained from 500 repetitions of the simulation. Vertical dashed lines delineate the events used for spike pattern analysis in (D). (**C**) Spike raster from all repetitions of the simulation. The “stim” arrow denotes the onset of the simulated thalamic inputs. (D) Distribution of spike patterns obtained in the simulation of a LIF neuron (inset: PSTH). (**E**) Distribution od spike patterns and PSTH (inset) for a model with modified parameters. Increasing the pre-synaptic firing rates of intracortical connections leads to higher coincidence of 101 and 011 patterns. In panels (A)-(D) the following parameters were used: *w*_exc_=0.0072 *μ*S, *τ*_exc_=0.9 ms, *f*_exc_=10 Hz, *w*_inh_=0.02252 *μ*S, *w*_Th_=0.035 *μ*S, *n*_Th_=28, *f*_Th_=700 Hz, *U*=0.65. In panel (E) four parameters were modified from this baseline: *f*_exc_=30 Hz, *w*_Th_=0.05*μ*S, *f*_Th_=300 Hz, *U*=0.7. All definitions and values of the remaining parameters are listed in Table 1.

We calculated the PSTH of the model by summing spike responses of *n* = 500 repetitions of the simulation. In each repetition the intracortical excitatory and inhibitory inputs, as well as the thalamocortical inputs, were drawn randomly from the Poisson distribution. In spite of this randomness, the model PSTH is composed of discrete peaks well separated by short valleys showing that the neuron fired precisely at preferred latencies (Figure 6B). Although the LIF model does not contain an explicit refractoriness, the intervals between the PSTH peaks correspond to the time required to depolarise the membrane from the reset potential (*V*_reset_ = −70 mV) to the spiking threshold (*V*_thr_ = −40 mV). In Figure 6A this time is seen as the slow rise time following the rapid downstrokes (reset) of the membrane potential triggered by spikes. Such a hyperpolarised period acts effectively as the refractory period as seen in the STPM and GLM.

The characteristic decay of the response in the somatosensory cortex observed in the experimental data (Figure 1B) could be driven by the adaptation of the neuron to the intense stimulation either at synaptic (Markram & Tsodyks, 1996) or neuronal level (Benda & Herz, 2003). Here, we model this process by means of short-term synaptic depression, which reflects the depression of thalamocortical synapses due to prolonged activity (Gil et al., 1997). The gradual decrease of synaptic drive makes the subsequent peaks smaller, broader, and separated by longer intervals (Figure 6B) as observed also in the experimental PSTH (Figure 1B). After 10 ms of stimulation the thalamocortical synapses deplete, abolishing further discharges.

In practice, the inputs to somatosensory cortex can also decay after a brief median nerve simulation (0.2 ms) applied to the median nerve invalidating our assumption of sustained synaptic drive. However, it has been found that the thalamocortical projections can respond with prolonged firing to brief presentation of the stimulus (Swadlow & Gusev, 2001). Interestingly, such responses also formed bursts of action potentials. If the axonal delays of multiple thalamocortical neurons are matched at the submillisecond level, such bursts could provide oscillatory inputs cortical level. The effects of such input patterns on the cortical responses should be investigated in the future.

The responses of the LIF model neuron vary from trial to trial (Figure 6C-D). This variability results from random cortical and thalamocortical inputs, which provide Poisson-distributed input spikes. Increasing the rate of inhibitory and excitatory inputs in a balanced fashion puts the neuron in a so-called noise-driven regime in which spikes are evoked by the random fluctuations over the threshold rather than by the mean depolarisation (Destexhe et al., 2003; Zerlaut et al., 2016). In this regime the responses of the neuron are more variable such that a broader range of different spike patterns is obtained across the trials. In particular, we found that the patterns with long latencies (such as, 011) or spike omissions (101) became more frequent at higher intracortical firing rates (Figure 6E).

In summary, the LIF model indicates that bursts in the somatosensory cortex can be driven by the input and do not always require intrinsic bursting mechanisms (reviewed by Krahe & Gabbiani (2004)). The number of spikes per burst and the within-burst intervals can be mechanistically explained by the integrating properties of single neurons that are equipped with an intrinsic adaptation process or driven by synapses that show short-term depression. Strong thalamic inputs can produce precise population responses at preferred latencies, which can overcome the variability. At the single-trial level the variability of the thalamic input is expressed in the form of stereotyped spike patterns.

## 4 Discussion

By means of simplified phenomenological models and a biophysical point-neuron model, we showed that within-burst variability of cortical S1 neurons can be decomposed into the private variability of each neuron and multiplicative input modulation that is shared by the entire population. The private variability explains most of the differences between responses elicited in single trials and underlies the re-appearance of the same spike patterns over multiple repetitions of the stimulus. The shared gain modulation coordinates the responses of many responding neurons and explains the puzzling co-variability between single-neuron and macroscopic population responses demonstrated in experimental recordings (Telenczuk et al., 2011). The models shed also light on the mechanism of S1 burst generation, their synchronisation across neurons, and suggest that spike patterns may encode time-varying cortical state at fast temporal scales.

### Mechanism of bursting

By means of a simple phenomenological model, the STPM, we showed that bursting in the primary somatosensory cortex results from the combination of intense synaptic bombardment and a refractory period. Such fast bursting triggered and sustained by an intense synaptic input has been termed “forced bursting” (Izhikevich, 2006).

The shape of a fitted recovery function in both models agrees well with the contribution of an afterhyperpolarisation (AHP) mediated potassium current and an afterdepolarisation (ADP) mediated either by persistent (Brumberg et al., 2000; Bal & McCormick, 1996) sodium or low-threshold calcium current (Jahnsen & Llinás, 1984): The initial dip, which we interpret as refractoriness, might reflect the AHP and the inactivation of sodium channels, whereas the subsequent over-shoot might correspond to the ADP. We note, however, that the over-shoot might be an artifact due to trial-to-trial variability (Figure 4C). We also demonstrated in a toy model that the over-shoot is not critical for bursting responses – the absolute refractory period combined with intense but transient inputs is sufficient to produce bursts with similar (but not identical) statistics (Figure 1C).

The STPM could also account for the correlations between interspike intervals (serial correlations). However, in a few neurons we found serial correlations differing from the ones it predicted. Since in these neurons processes occurring at long time scales could shape the spike patterns, we fitted them with the GLM, which considers spike-history effects extending to multiple interspike intervals. We found that the GLM with a horizon of 8 ms provided an optimal fit to these data in agreement with the time scales of short-term synaptic plasticity (Tsodyks & Markram, 1997) and firing-rate adaptation (Benda & Herz, 2003). The latter is often mediated by the slow AHP currents providing another link between a biophysical process and the recovery function of our phenomenological model.

The significant serial correlations could be also explained by a model which includes trialto-trial variations of the input intensity (gain modulation). We found that introducing such variations in the model results in the over-estimation of the serial correlations estimated from the simulated data. In addition, these variations may lead to the over-estimation of the recovery function in form of the overshoot appearing briefly after the absolute refractory period. Although such an overshoot is also present in the recovery function estimated from the data, we believe that it is not an artifact of the estimation method. First, the modulation must be strong (*γ* = 0.8) to produce a visible overshoot, whereas we found that modest modulation (*γ* = 0.2) is consistent with the serial correlation and EEG correlation estimated in the data. Secondly, we found that GLM is robust with respect to such modulation introduced in the simulated model, but still it uncovers an overshoot in the experimental data. Nevertheless, in the future it would be instructive to extend the spike-train models (STPM and GLM) with the fluctuating gain factor and fit it directly to the data.

We were able to reproduce qualitatively both the average and single-trial features of the burst responses in a more realistic leaky integrate-and-fire neuron. Although such models are a gross simplification of the real neurons both in terms of spiking mechanism and morphological features, it has been suggested that the LIF may faithfully reproduce some features of spike generation (Brette, 2015). In the model, the within-burst interval was controlled by the time required to reach the threshold from the hyperpolarised state (membrane time constant), and the gradual decay of the amplitude of PSTH peaks was due to the short-term synaptic depression. The latter mechanism can be related to the depletion of the available vesicles in the pre-synaptic terminal (Markram & Tsodyks, 1996). However, it would be possible to replace it with some other form of adaptation (Brette & Gerstner, 2005). Both mechanisms lead to extinction of the initial synaptic drive, which explains the burst-like transient response to the step-like thalamic inputs. We note, however, that without recordings from thalamocortical projection neurons we can not infer the inputs of the cortical neurons. Our models are still compatible with temporally structured inputs.

The trial-to-trial variability of the model was due to variable arrival times of the thalamic inputs, but also due to the intra-cortical inputs. The latter were configured such that the neuron was in the “high-conductance state” reproducing the property of cortical neurons receiving constant bombardment of inhibitory and excitatory inputs (Destexhe et al., 2003). Apart from decreasing the membrane time constant thus allowing for rapid repeated discharges, these intracortical inputs introduced substantial trial-to-trial variability that could explain the observed spike pattern distribution.

Previous studies have shown that most of the bursting neurons in the S1 macaque cortex are characterised by broad spikes, which suggests that they are pyramidal neurons or spiny stellate neurons (Baker et al., 2003). This is confirmed by intracellular recordings in barrel cortex showing that regular spiking neurons but not intrinsic bursting neurons followed the phase of high-frequency oscillations in surface recordings (Jones et al., 2000). Our results are consistent with these findings and strengthen the evidence that a subclass of S1 neurons activated by median nerve stimulation belongs to the regular spiking neurons. However, a subset of neurons analysed here (5 of 17 neurons) did also fire bursts that were not locked to the median nerve stimulation showing that at least some of them may belong to the intrinsic bursting class.

### Burst synchronisation

A striking feature of the S1 bursting is that the signature of the burst also appears in macroscopic signals such as the EEG. The visibility of the burst in the surface recordings was interpreted as a sign of strong synchronisation between the neurons (Jones et al., 2000), which could be mediated, for example, by fast synaptic potentials or gap junctions (Draguhn et al., 1998). By extending our model to a population of uncoupled neurons, we demonstrated that the sub-millisecond synchronisation between multiple neurons does not require a fast coupling mechanisms, but results from shared synaptic inputs arriving through thalamocortical fibers. Provided that the biophysical properties of the receiving population and axonal conduction times vary in a narrow range, these inputs will elicit synchronous bursts of spikes. The required precision in the arrivals of afferent spikes could be achieved by means of a plasticity rule that selects inputs arriving synchronously at the cortical synapses (Gerstner et al., 1996).

### Role of spike patterns

Trial-to-trial variations in S1 responses can be classified into a set of spike patterns defined by the occurrences of spikes within 10-ms-long bursts (Telenczuk et al., 2011). Such temporal patterns of neuronal responses were first identified in cat striate cortex and crayfish claw (Dayhoff & Gerstein, 1983), and later in the temporal cortex of monkeys, cat lateral geniculate nucleus (Fellous et al., 2004) and in the rat hippocampus (Diba & Buzsaki, 2007; Schmidt et al., 2009).

Here, we proposed a model in which the occurrence of spike patterns is regulated by the input intensity, that is the rate of incoming spikes; in contrast precise timing of the input was not necessary. The temporal information stored in the spike patterns is complementary to the output rate (spike count) in the sense that the spike patterns with identical number of spikes (and therefore the same output rate) could still provide extra information concerning its inputs. For example, the early (110) doublet is more common for high input intensity; the opposite is true for the late (011) doublet (Figure 5B). This mechanism could be especially useful for encoding inputs that would normally exceed the maximum firing rate set by the refractory period.

In one study the stimulus intensity was related to the within-burst intervals of spike responses recorded in the dorsal lateral geniculate nucleus (dLGN) (Funke & Kerscher, 2000). Our results are consistent with this hypothesis. In the STPM, the within-burst intervals are constrained by the refractory period, but their length can also vary as a function of the synaptic drive (intensity function). In addition, the length of refractory period may not be fixed but it might be modulated by the firing rate. It has been shown that models allowing for this modulation may better describe the spike times in response to the time-varying stimulation (Koyama & Kass, 2008).

Short trains of spikes are also well suited to evoke specific synaptic response or trigger synaptic plasticity (Lisman, 1997; Song et al., 2000; Swadlow & Gusev, 2001; Tsodyks & Markram, 1997; Maass & Zador, 1999), they are optimally placed to represent neuronal variables in a form that is easily processed, stored and transmitted (Leibold et al., 2008; Tiesinga et al., 2008). In this spike-timing-based view, neural systems take advantage of the temporal information encoded into spike patterns to represent slowly-changing cortical states (such as attention or waking). Alternatively, spike patterns could also allow for more reliable representation of neuronal inputs (Toups et al., 2012). These rate-based and spike-timing-based interpretations of spike patterns are not contradictory and could even act as independent communication channels (Tiesinga et al., 2008).

We showed that the distribution of spike patterns over neurons and the amplitude of the averaged population signal are regulated by input magnitude, which could reflect gating of neuronal signals through attention, expectation, sleep and waking (Fontanini & Katz, 2008; Shu et al., 2003; Steriade et al., 2001). Similar gain control mechanisms were implemented in realistic neural models through, for example, concurrent modulation of excitation and inhibition (Hô & Destexhe, 2000; Chance et al., 2002; Vogels & Abbott, 2009) or short-term synaptic depression (Rothman et al., 2009). More generally, multiplicative noise can account for the variability and co-variability of neuronal responses in the thalamus and many cortical areas, including the lateral geniculate nucleus, V1, V2 and MT (Goris et al., 2014).

### Population correlates of spike patterns

Macroscopic signatures of the bursts were shown to match the somatosensory-evoked potentials in monkey epidural EEG and human scalp EEG, so the high-frequency EEG burst might link the non-invasive macroscopic recordings and microscopic neuronal activity (Curio, 2000; Telenczuk et al., 2011). We could reproduce this puzzling relation between the single-neuron spike patterns and the macroscopic EEG signals by means of the STPM with the gain modulation. To model the high-frequency EEG signals, we used the high-pass filtered average response of the population (population PSTH). This choice was motivated by previous studies on the origins of the high-frequency EEG signals: while the low-frequency field potentials are known to correlate mostly with the synaptic currents (Buzsáki et al., 2012; Mazzoni et al., 2015), it has been recently demonstrated that the population spike rate is a better predictor of high-frequency >400 Hz EEG power (Telenczuk et al., 2015, 2011). Based on these results we conclude that the link between the microscopic and macroscopic activity could be partially explained by the neuronal correlations mediated by the common gain modulation.

To sum up, our modelling shows how the characteristic features of the spike burst, i.e., its frequency and amplitude, can be related to the biophysical properties of neurons, such as refractory period and short-term synaptic depression, whereas the internal burst composition is controlled by the background activity and gain modulation. As a conclusion we argue that the brain could use small within-burst timing differences to encode the dynamical cortical state at time scales amenable to neuronal processing.

